# In-silico screening of drug candidates for thermoresponsive liposome formulations

**DOI:** 10.1101/2020.05.11.087742

**Authors:** Martin Balouch, Martin Šrejber, Marek Šoltys, Petra Janská, František Štěpánek, Karel Berka

**Affiliations:** Department of Chemical Engineering, University of Chemistry and Technology, Prague, Technická 3, 166 28 Prague 6, Czech Republic; Regional Centre of Advanced Technologies and Materials, Department of Physical Chemistry, Faculty of Science, Palacký University Olomouc, 17. listopadu 12, 771 46 Olomouc, Czech Republic

## Abstract

Liposomal formulations can be advantageous in a number of scenarios such as targeted delivery to reduce the systemic toxicity of highly potent Active Pharmaceutical Ingredients (APIs), to increase drug bioavailability by prolonging systemic circulation, to protect labile APIs from degradation in the gastrointestinal tract, or to improve skin permeation in dermal delivery. However, not all APIs are suitable for encapsulation in liposomes. Some of the issues are too high permeability of the API across the lipid bilayer, which may lead to premature leakage, too low permeability, which may hinder the drug release process, or too strong membrane affinity, which may reduce the overall efficacy of drug release from liposomes. Since the most reliable way to test API encapsulation and release from liposomes so far has been experimental, an *in silico* model capable of predicting API transport across the lipid bilayer might accelerate formulation development. In this work, we demonstrate a new *in silico* approach to compute the temperature dependent permeability of a set of compounds across the bilayer of virtual liposomes constructed by molecular dynamics simulation. To validate this approach, we have conducted a series of experiments confirming the model predictions using a homologous series of fluorescent dyes. Based on the performance of individual molecules, we have defined a set of selection criteria for identifying compatible APIs for stable encapsulation and thermally controlled release from liposomes. To further demonstrate the *in silico*-based methodology, we have screened the DrugBank database, identified potent drugs suitable for liposome encapsulation and successfully carried out the loading and thermal release of one of them - an antimicrobial compound cycloserine.

## 1 Introduction

Liposomes are spherical vesicles containing an internal aqueous cavity enclosed by a bilayer usually made of phospholipids, often with the addition of sterols. At low temperatures, the lipid bilayer exists in an ordered state and is non-permeable for hydrophilic payloads; the content of the internal cavity is trapped inside and protected from the outside environment ^1^. Therefore, liposomes can be used as carriers for the encapsulation and controlled release of active substances including small-molecule drugs ^2^, biomolecules ^3^ and genes ^4, 5^), or as building blocks for more complex drug delivery systems ^6^. Beyond pharmaceutics, liposomes find applications in cosmetic ^7^ or nutraceutical ^8^ formulations. To this day, liposomes were successfully used in several FDA approved products available on the market ^9^.

The properties of liposomes can be tailored to specific delivery and release mechanisms ^8, 10–12^. Among the most critical parameters affecting the storage stability and the final application properties of liposomes are the bilayer composition, lamellarity and size ^1, 13, 14^. These parameters can be influenced by the liposome preparation method, which typically involves the dissolution of lipids in an organic solvent and subsequent dispersion into aqueous media ^15^ with or without relying on membrane permeation of the encapsulated solute ^16^. The solute permeation mechanism at the point of use can include spontaneous loss of lipid bilayer integrity upon interaction with living cells ^17^ or an externally triggered event such as local hyperthermia ^10^. Depending on its composition, the lipid bilayer has a phase transition temperature at which it changes from a highly ordered into a disordered state where it becomes significantly more permeable for the encapsulated solutes ^18^. This transition between states is usually reversible ^19^, and can be triggered by incorporated magnetic nanoparticles ^20^ stimulated by an external magnetic field. This offers interesting possibilities for customised controlled release scenarios, e.g. in theranostics.

However, the actual permeability through the lipid bilayer is specific to the encapsulated compound and strongly depends on the liposome composition. The encapsulated compound needs to be able to permeate through the bilayer to escape the liposome when heated, but it should not permeate when the lipidic bilayer is in the ordered state. The occurrence of the following cases makes a compound incompatible with a given liposome composition: *(i)* it is permeable not only in the disordered state but also in the ordered state, resulting in premature leakage; *(ii)* a strong repulsion between the bilayer and the compound will prevent permeation even in the disordered state; *(iii)* a strong attraction towards the bilayer will not permit the compound to be released. Predicting these behaviours for specific compounds in combination with a particular lipid bilayer composition and its physical state is a non-trivial task and exploring all the possible compositions experimentally would require an extensive parametric study. Formulation optimisation by trial-and-error is therefore extremely time consuming and often leads to the rejection of the entire liposome formulation route for a given compound simply because a limited number of experiments performed within a random sub-region of the parametric design space did not result in a “hit”.

Recently, *in silico* molecular modelling became increasingly feasible as an accurate predictive tool, which can reduce the need for extensive experimental parametric studies during formulation development ^21, 22^. The availability of computational power in combination with efficient algorithms made it possible to construct atomistic models of lipid bilayers based on molecular dynamics simulations ^23, 24^, which in turn makes *in silico* models a realistic alternative to physical experiments. Hence, the aim of the present work was to develop and validate an *in silico* tool specifically for the case of encapsulation and thermally induced release of small-molecule payloads from liposomes. The proposed strategy was to utilise a computational model ^25, 26^ to construct a virtual three-component phospholipid bilayer in different conditions and states (i.e. below and above its phase transition temperature) using molecular dynamics, and then to predict the permeability of a set of compounds through the bilayer using quantum chemistry based statistical thermodynamic computation. The calculated predictions were then validated by conducting physical experiments using liposomes loaded with the same compounds – a homologous series of fluoresceins. Once the model was experimentally validated, a series of potent drug molecules from the DrugBank database ^27^ were analysed by the computational model in order to predict which of them might be suitable for liposome encapsulation and temperature-responsive release. Based on the *in silico* selection, an antimicrobial compound cycloserine was successfully formulated in liposomes and its thermo-responsive release was demonstrated experimentally.

## 2 Methods

### 2.1 Computational methods

#### 2.1.1 Molecular dynamics simulations of lipid bilayers

All molecular dynamic (MD) simulations were performed using the GROMACS 4.5.4 software package ^28^. Model of a membrane mixture containing premixed 1,2-dipalmitoyl-sn-glycero-3-phosphocholine (DPPC): 1,2-dipalmitoyl-sn-glycero-3-phosphoglycerol (DPPG): cholesterol (Chol) at a molar ratio of 75:10:15 was created using an in-house script with a total number of 128 lipids (64 lipids per leaflet). For simulations of the lipid bilayer the Slipids parameters ^29^ were used along with TIP3P water model ^30^. Na^+^ and Cl^−^ counterions were added to maintain electroneutrality of the system and to reach physiological concentration. Each bilayer system was simulated for 250+ ns until equilibrium was reached (see area of lipid evolution at Supporting Information Figure S1.1) The temperature was kept constant using the Nosé-Hoover thermostat with a coupling constant of 0.5 ps ^31, 32^). The pressure was kept constant at 1 atm by Parrinello-Rahman barostat and semi-isotropic coupling with a coupling constant of 10 ps and isothermal compressibility of 4.5 × 10^−5^ bar^−1 33^. The long-range electrostatic interactions were calculated using the particle-mesh Ewald scheme with a 1.6 nm cutoff ^34^. The cutoff for Lennard-Jones interactions was set to 1.0 nm with switching the function to zero at 1.5 nm. The LINCS algorithm was used to constrain all bonds ^35^. Periodic boundary conditions were applied in all directions. The temperature dependence of membrane model was simulated in the range between 293 K and 333 K (Figure 1).

**Figure 1.**
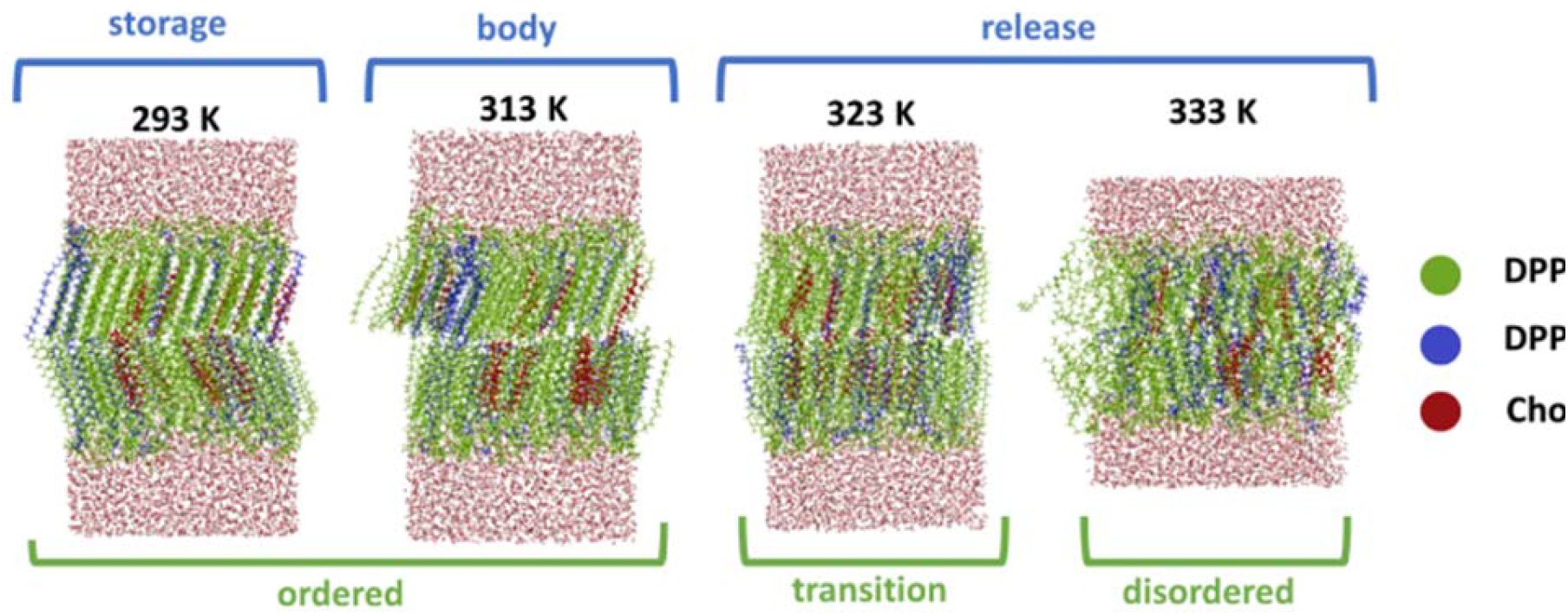
Structures of mixed DPPC:DPPG:Chol (75:10:15) membranes obtained by MD simulations at various temperatures corresponding to liposome storage, human body temperature and release of compounds from liposomes.

#### 2.1.2 COSMOmic/COSMOperm calculations of solute permeation

The LigPrep and MacroModel packages ^36, 37^ were used to generate neutral conformers of compounds in vacuum using the OPLS_2005 force field ^38^. For each compound a maximum of 10 conformers (within 5 kcal/mol of conformer with the lowest energy and RMSD at least 0.2 nm between individual conformers) were selected based on MCMM/LMC2 conformation searching algorithm with Monte Carlo structure selection. For each conformer, a subsequent DFT/B-P/cc-TZVP vacuum and COSMO optimisation were carried out using Turbomole 6.3 ^39^ implemented in cuby4 framework ^40^ followed by a single-point energy DFT/B-P/cc-TZVP calculation with *fine grid* option to obtain the COSMO files. The overall .mic files for lipid bilayers were obtained by fitting individual lipid COSMO files to an average of 10 bilayer structures from the MD simulations. For each compound, conformer/lipid bilayer system COSMOmic/COSMOperm X18 calculations ^41, 42^ were performed to obtain averaged free energy profiles and hence the partition (logK_lip/wat_) and permeability (logP_erm_) coefficients. The procedure of these calculations is described in Supporting information. Initially, a set of 5 fluorescent chemical compounds was used for validation purposes: fluorescein, 5-carboxyfluorescein, 6-carboxyfluorescein, fluorescein-5-isothiocyanate, and calcein. Once the computational model was validated experimentally (Section 2.2), the permeation of a further 56 compounds selected from the DrugBank database was computed and used as a basis for proposing criteria for drug selection with respect to their suitability for liposome encapsulation and release. These 56 compounds were selected based on their high toxicity on living organisms and therefore potential benefits of a liposomal formulation. A full list of the compounds is provided in Section 3.4. Calculations of all molecules were performed 10 times. The most promising drug candidate was then selected based on its logK_lip/wat_ and logP_erm_ values and its liposomal formulation and thermoresponsive release was demonstrated experimentally (Section 2.3).

### 2.2 Experimental validation of model predictions

#### 2.2.1 Materials

Cholesterol (Chol, >99%), ammonium hydroxide solution (28–30%), phosphate-buffered saline (PBS) tablets, 5-(6)-carboxyfluorescein (5(6)-CF, 95%), calcein (C, >95%), D-cycloserine, fluorescein (F, 95%), fluorescein-5-isothiocyanate (FITC, 90%) and TRITON® X-100 were purchased from Sigma-Aldrich. Ethanol (for UV spectroscopy, 99.8%), chloroform (p.a.), sodium chloride (>99%), sodium hydroxide (p.a.), methanol (p.a.) and hydrochloric acid (HCl, 35%, p.a.) were purchased from Penta. Dipalmitoylphosphatidyl-choline (DPPC) and sodium dipalmitoylphosphoglycerol (DPPG) were purchased from Corden Pharma. All chemicals were used as received. Deionized water (Aqual 25, 0.07 μS/cm) was used for all reactions and treatment processes.

#### 2.2.2 Preparation of liposomes

Liposomes were prepared using the Bangham method. Briefly: 10 mg of lipid and cholesterol mixture was dissolved in 2 mL of chloroform: methanol mixture (2:1) in a 25mL round flask. The mixture was evaporated in a rotary evaporator at a constant temperature of 55 °C and gradually lowering pressure from atmospheric to 150 mbar. The dried lipids formed a film around the flask. The sample was then kept under vacuum for at least 12 hours. For further rehydration, 2 mL of a hydration phosphate buffer solution containing the substance to be encapsulated was added to the flask (the compounds and their concentrations are listed in Table 1). The content of the flask was then heated and stirred using a vortex shaker until no visible lipid film pieces were present on the flask walls. 1 mL of the obtained lipid suspension was then transferred to a liposome extrusion device (Avanti Polar Lipids), heated to 69 °C and extruded through a porous membrane with 800 nm pores 21 times to decrease and unify the size of the formed liposomes. The size distribution of the prepared liposomes was measured using the Malvern Zetasizer. The liposomes wre also visualised using transmission electron microscopy (TEM, Jeol JEM-1010, acceleration voltage 80 kV).

**Table 1.** Solutions used for the lipid film hydration

| Fluorescent dye | Abbreviation | Concentration<br>[mg/mL] | pH |
| --- | --- | --- | --- |
| Calcein | Cal | 5.0 | 7.4 |
| 5(6)-Carboxyfluorescein | 5(6)-CF | 7.5 | 7.4 |
| Fluorescein | F | 2.5 | 7.4 |
| Fluorescein-5-isothiocyanate | FITC | 0.1 | 7.4 |

#### 2.2.3 Measurement of thermo-responsive release

For experimental validation of the computational model, fluorescein derivatives with systematically varying chemical properties have been chosen (Table 1). Since the release measurement is based on the fluorescence emission of the dye dissolved in an aqueous supernatant, only water-soluble fluorescent dyes were used. Four fluorescein derivatives were found to be sufficiently soluble for this purpose. From these four compounds, near-saturated solutions in water were made and used for hydration of the lipid film to form liposomes. The actual concentrations and pH values used for liposome encapsulation are summarised in Table 1.

The measurement of thermally induced release of these substances from the liposomes was based on repeated liposome separation by centrifugation and fluorescence emission intensity measurement of the supernatant (Cary Eclipse Fluorescent Spectrophotometer, Agilent). First, four cycles of centrifugation, supernatant removal, dilution and redispersion were conducted to remove any non-encapsulated dye present in the samples after liposome preparation. After liposome extrusion, approximately 1 mL of the liposome dispersion was transferred to several 2mL Eppendorf vials and cooled to room temperature. The samples were then filled up to the to 2mL mark with a buffer solution of the same ionic strength as the hydration solution. Then, centrifugation at 5000 RCF (Minispin, Eppendorf AG Hamburg) for 15 minutes was performed. The supernatant was removed and replaced with a fresh buffer solution, the liposomes were redispersed and again centrifuged. After the last centrifugation, the fluorescence emission intensity of the supernatant was measured to verify the completion of the purification. The excitation wavelength was set to 490 nm and the emission spectra were measured at the maximum intensity for each dye (between 515 nm and 525 nm). Based on the calibration using the same settings, concentration of dye was calculated. From the difference of measured dye quantities before and after release, the amount of released dye was calculated.

To determine the total quantity of the encapsulated dye inside the liposomes, 2 μL of TRITON® X-100 was added to the sample, which resulted in membrane destabilisation and release of all the dye that was present either in the liposomal cavity or in the membrane. In order to measure thermally induced release from liposomes, the purified samples were heated to 65 °C for 1 hour to bring the liposome membranes into the disordered state and make them more permeable. In both measurements (membrane destruction using TRITON® and thermal release), the samples were centrifuged once more and the fluorescence emission intensity of the supernatant was measured. From the difference of the fluorescence emission before and after TRITON® addition or heating, the released quantity of the fluorescent dye was calculated.

Since each of the four fluorescent dyes was used at different concentration (due to difference in their aqueous solubility), a theoretical loading capacity for each dye was calculated in order to allow effective comparison of the samples. For simplicity, it was considered that all phospholipids used in the experiment were DPPC and the liposome size was uniform at 800 nm in diameter. The area occupied by a single DPPC lipid was considered to be 0.6 nm^2^. ^43^ From the known area of one lipid and the known surface area of one liposome, the quantity of DPPC molecules forming one liposome was calculated. With the known mass of lipids in one experiment (10 mg), the molar mass of DPPC (734 g/mol) and the concentration of fluorescent dye in the hydration solution, the theoretical loading capacity could then be obtained:

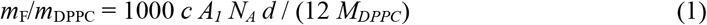

where *m*_F_/*m*_DPPC_ is the theoretical capacity in μg of solute per 1 mg of lipids, *c* is the fluorescent dye concentration in mg/mL, *A*_1_ is the area of one DPPC molecule, *d* is the mean liposome diameter, *N*_*A*_ is the Avogadro constant and *M*_DPPC_ is the molar mass of DPPC (734 g/mol). In this theoretical calculation, it was assumed that the hydration solution was encapsulated inside all liposome cavities. Upon release, the entire volume of the encapsulated solution and the corresponding quantity of the dissolved dye was considered as the theoretical maximum capacity of the liposomes prepared.

For the possibility of comparison between experiments and calculations, which were made for unionized species, the fraction of unionized species was calculated using MarvinSketch 20.16. ^44^

#### 2.2.4 Visualisation of membrane partition and release

Laser scanning confocal microscopy (Olympus Fluoview FV1000 with Canon EOS 100D camera) was used to visualise the liposomes and their surrounding medium both before and after the thermally induced release of the encapsulated dyes. Furthermore, liposomes loaded with all four used dyes, centrifuged and redispersed were also visualised. The prepared liposomes were not extruded through the 800 nm membrane after in order to keep their size large (for easier observation). For the confocal images Olympus UPLSAPO Objective 40x, excitation wavelength 405 nm and fluorescence detection 505-540 nm were used.

### 2.3 Liposomal formulation of drug identified by *in silico* model

The most promising substance identified by the *in silico* model from a sample of 56 entries in the DrugBank database (cycloserine) was encapsulated into liposomes as described in Section 2.2. The stability of liposomes at storage temperature and the ability to release the substance above the phase transition temperature of the lipid bilayer was demonstrated by measuring the proliferation of bacteria *E.coli* using the resazurin assay.^45^ The bacteria suspension was grown overnight in LB media at 37 °C, shaken at 200 rpm. The inoculate was diluted to the appropriate cell density (OD = 0.06) and seeded on 96-well plates. The assay composed of a row of samples exposed to liposomes before release, liposomes after heat release (60 °C for 30 minutes), liposomes after membrane disruption by TRITON® as a positive control, and a row of bacteria with added antibiotic (50 μg/mL kanamycin) as a further positive control. Each well was filled with 100 μl of the sample (liposomes loaded with 10 mg/mL cycloserine, repeatedely diluted and centrifugated). Then, 20 μl of diluted bacterial suspension was added into all wells and mixed thoroughly. Samples after release and antibiotics were performed in triplicates and sample before the release in duplicates of both samples, which were later used for release. After overnight incubation at 37 °C, 20 μL resazurin was added to all wells and incubated at 37 °C for another 4 h. Changes in colour were observed. Fluorescence measurements were performed in a spectrofluorometer (Infinite 200 PRO, Tecan Austria) at 560 nm excitation and 590 nm emission wavelength. Bacteria without any antibiotics were used as a negative control.

## 3 Results

### 3.1 Computational studies

#### 3.1.1 MD simulations of bilayer state at various temperatures

Molecular dynamics simulations of the mixed DPPC:DPPG:Chol (75:10:15 molar ratio – chosen from previous experimental experiences as an ideal composition for encapsulation of 5(6)-CF [10]) membrane model were carried out at temperatures corresponding to the liposome storage temperature (293 K), the body temperature (313 K) and the temperatures presumed to be required for the release of encapsulated solute (323 K and 333 K). At lower temperatures for the liposome storage, the membranes were found to be in a highly ordered phase (Figure 1). An increase of temperature by 20 K to the body temperature did not result in any changes in the phase ordering, and the membrane remained in an ordered phase. The dependent simulations showed a sharp change in the membrane structure between 323 K and 333 K, where a phase shift into the disordered phase was observed. This shift is located slightly higher than experimentally observed phase transition temperature of this membrane mixture [10] (315 K). Above 323 K the lipid bilayer formed a disordered structure preferable for the release of the encapsulated compounds from the internal liposomal cavity.

#### 3.1.2 COSMO-based partitioning and permeability calculations for fluorescent dyes

The values of temperature-dependent partition coefficients logK_lip/wat_ obtained from the COSMOmic calculations compared in Fig. 2A showed no significant changes with increasing temperature for the given fluorescent compounds. The lipophilic character of the compounds increased in the order of Cal < 6-CF < 5-CF < F < FITC but the drug partitioning between water and the lipidic membrane was not found to be much sensitive to temperature changes while all the compounds became only slightly more hydrophobic at elevated temperatures.^46^ In contrast, the permeability coefficients logP_erm_ obtained from COSMOperm calculations have shown a significant increase with the temperature of about two orders of magnitude within 40K difference and varied widely between the compounds (Fig. 2B). We have therefore analysed the performance of each dye in both partitioning and permeability at the lowest (293 K) and the highest (333 K) temperature in more detail.

**Figure 2.**
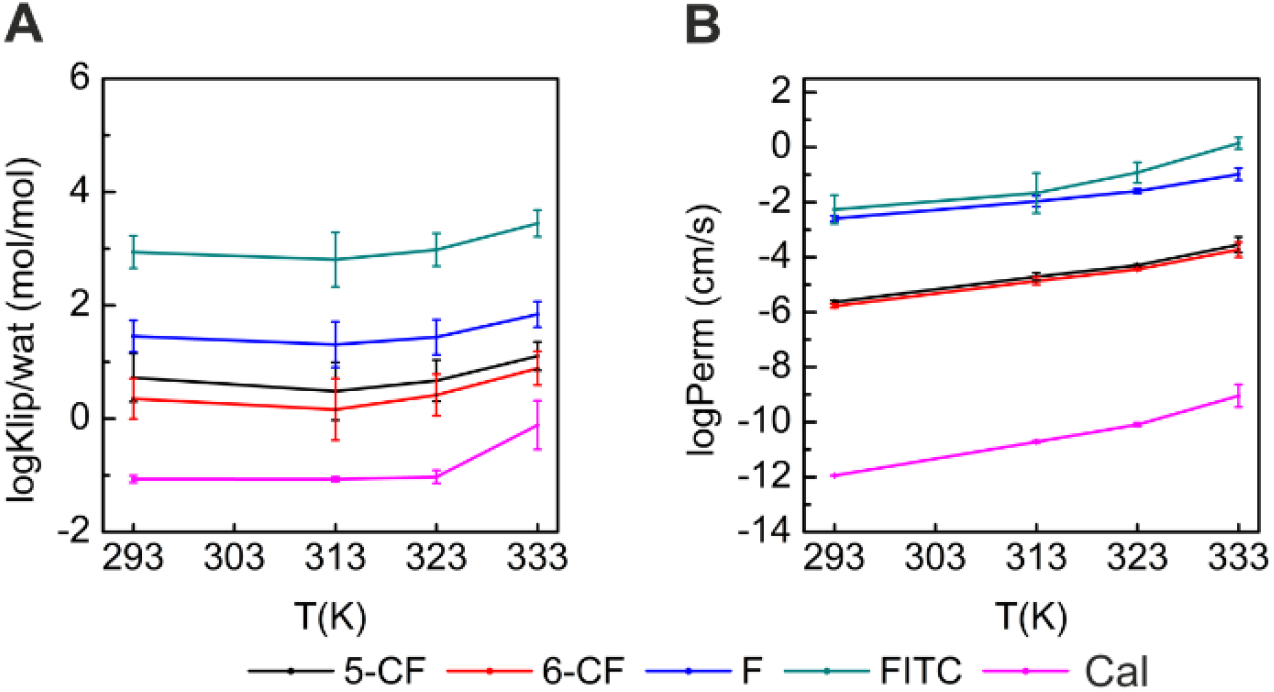
Computed temperature dependence of membrane/water partition coefficients logK_lip/wat_ [mol/mol] (panel A) and permeability coefficients logP_erm_ [cm/s] (panel B) for a series of five fluorescent dyes on the DPPPC:DPPG:Chol mixed membrane.

Each fluorescent derivative has shown a different behaviour in terms of the partitioning coefficient and permeability. The dye with the highest partition coefficient is fluorescein-5-isothiocyanate (FITC), having a logK_lip/wat_^293 K^ = 2.94 ± 0.28. This would suggest a strong presence of the dye within the membrane while only a small amount of FITC would remain encapsulated in the internal aqueous cavity of the liposome. At the same time, FITC would easily permeate through the membrane due to its high permeability coefficient (logP_erm_^293 K^ = −2.26 ± 0.52) even at low temperature. As such, FITC does seem suitable for encapsulation in membrane cavity but not for temperature-controlled release in the used lipid mixture. For fluorescein (F), a slightly lower partition coefficient logK_lip/wat_^293 K^ = 1.45 ± 0.28 would suggest a shift of equilibrium towards the aqueous phase. Therefore, it is likely that fluorescein would prefer equivalent partitioning into the aqueous core and the lipidic membrane. However, the high permeability at all temperatures (logP_erm_ = −2.59 ± 0.09, –1.60 ± 0.09 and –0.98 ± 0.22 at 273 K, 323 K and 333 K, respectively) again does not make it a promising candidate for liposome encapsulation.

Relatively low partition coefficients were observed for 5-carboxyfluorescein (5-CF) and 6-carboxyfluorescein (6-CF), namely logK_lip/wat_^293 K^ = 0.72 ± 0.43 and 0.35 ± 0.36, respectively. Therefore, these compounds should prefer the aqueous phase within the liposomal cavity. Moreover, their permeability is nicely temperature-dependent since they have a low permeability at low temperatures (logP_erm_^293 K^ = −5.64 ± 0.06 and −5.78 ± 0.06, respectively) and two orders of magnitude higher permeability at high temperatures (logP_erm_^333 K^ = –3.55 ± 0.28 and –3.73 ± 0.27, respectively). As a result, these two compounds appear to be suitable candidates that could provide stable liposomal encapsulation during storage, followed by temperature triggered release.

Finally, calcein (Cal) shows the lowest values for both partitioning and permeation coefficients. The lowest partitioning logK_lip/wat_^293 K^ = –1.07 ± 0.07 at all temperatures among all dyes suggests predominant encapsulation of the drug in the water phase within the liposomal cavity, which seems like a great property to block drug release from the storage liposome. However, calcein also shows an extremely low ability to permeate through the bilayer and leave the liposome even at elevated temperatures (logP_erm_^333 K^ = –9.05 ± 0.41). Therefore, calcein is probably a bad candidate for thermally controlled drug release, although it might be suitable for other release mechanisms.

These *in silico* predictions suggest that out of the tested set of fluorescent dyes, the most promising candidates for encapsulation and temperature-controlled release from DPPC:DPPG:Chol (75:10:15 molar ratio) liposomes are 5-CF and 6-CF, while FITC and F seem to be too permeable and Cal will not be able to permeate even at higher temperatures.

### 3.2 Experimental verification

#### 3.2.1 Liposome properties

To validate our theoretical predictions, we have prepared dye-loaded liposomes as described in Section 2.2. The size of the prepared liposomes was centred at 800 nm irrespective of the loaded dye (Fig. 3A). However, a fraction of liposomes with a size around 100 nm was present in all samples and also confirmed by TEM analysis (Fig. 3B). We suggest that the smaller liposomes (100 nm) do not settle during the centrifugation steps and are disposed of together with the supernatant during the purification process. This can be a possible source of losses of the encapsulated substance and thus somewhat lower measured quantities of the released dye, as discussed below.

**Figure 3.**
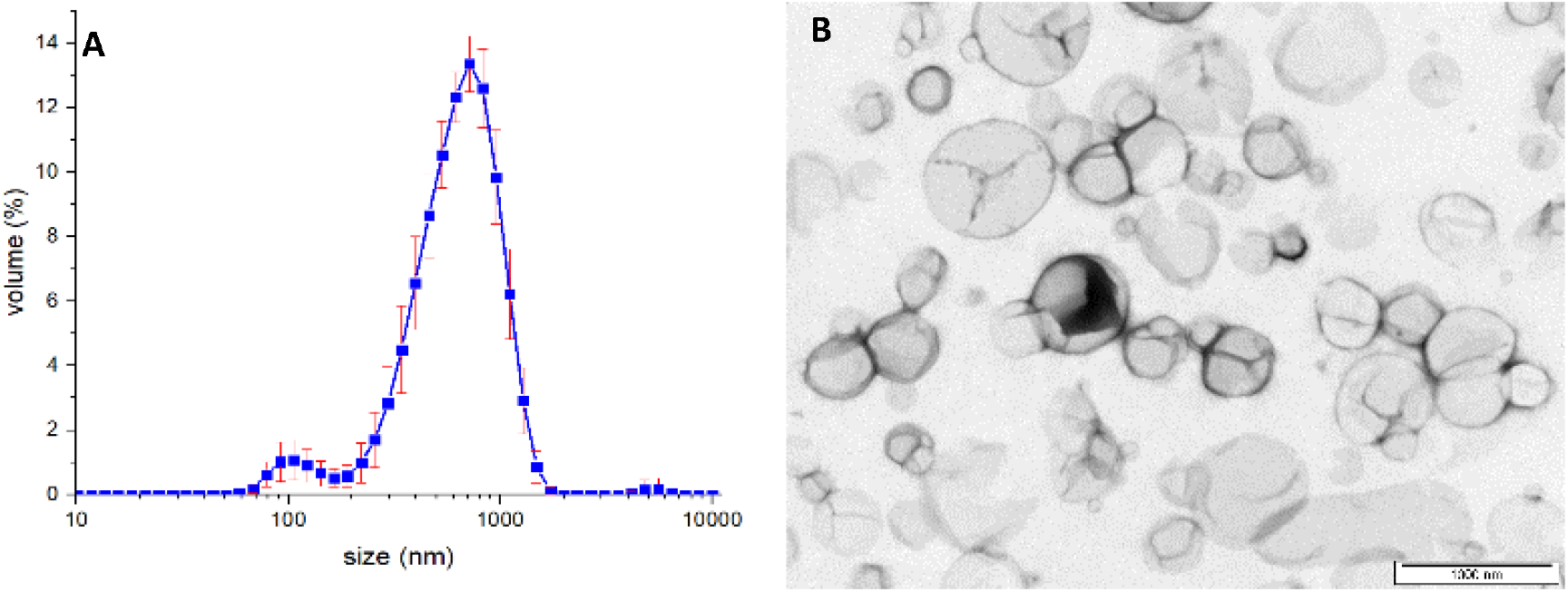
Volume size distribution (panel A) and TEM micrograph (panel B) of the prepared liposomes. The scale bar represents 1000 nm.

#### 3.2.2 Encapsulation and release experiments with fluorescent dyes

The loading of 5(6)-Carboxyfluorescein into liposomes was rather efficient, achieving 73 ± 35 μg_dye_/mg_lipid_, which represents approximately 30 % of the theoretical capacity (Table 2). The dye remained stable inside the liposomes when kept at the storage temperature (Fig. 4A), but it was readily released upon heating to 333 K (62 ± 8 μg_dye_/mg_lipid_). This shows that 5(6)-CF is a good example of a molecule to be used for encapsulation in liposomes with thermally controlled release, confirming the theoretical calculations. From the comparison between theoretical and experimental results we can conclude that logP_erm_ = −5.6 at 293 K is not high enough to allow premature membrane permeation. On the other hand, logP_erm_ = −3.6 at 333 K seems to be sufficient for release – this opens a window for applicability.

**Table 2.** Measured release from liposomes after heating

| Dye | pH<sup>†</sup> | Released after heating<br>($\mu\text{g}_{\text{dye}}/\text{mg}_{\text{lipid}}$) | Released after membrane disruption<br>($\mu\text{g}_{\text{dye}}/\text{mg}_{\text{lipid}}$) | Theoretical capacity<br>($\mu\text{g}_{\text{dye}}/\text{mg}_{\text{lipid}}$) | Fraction of unionized species<br>(%) |
| --- | --- | --- | --- | --- | --- |
| 5(6)-CF | 7.4 | $62 \pm 8$ | $73 \pm 35$ | 246 | 0.03 % |
| F | 7.4 | 0.0 | $0.3 \pm 0.1$ | 82 | 95.4 % |
| FITC | 7.4 | 0.0 | $0.6 \pm 0.2$ | 3.3 | 95.4 % |
| Cal | 7.4 | $0.03 \pm 0.02$ | $0.4 \pm 0.1$ | 164 | < 0.01 |

<sup>†</sup> hydrating solution: 10 mM PBS adjusted to pH 7.4

**Figure 4.**
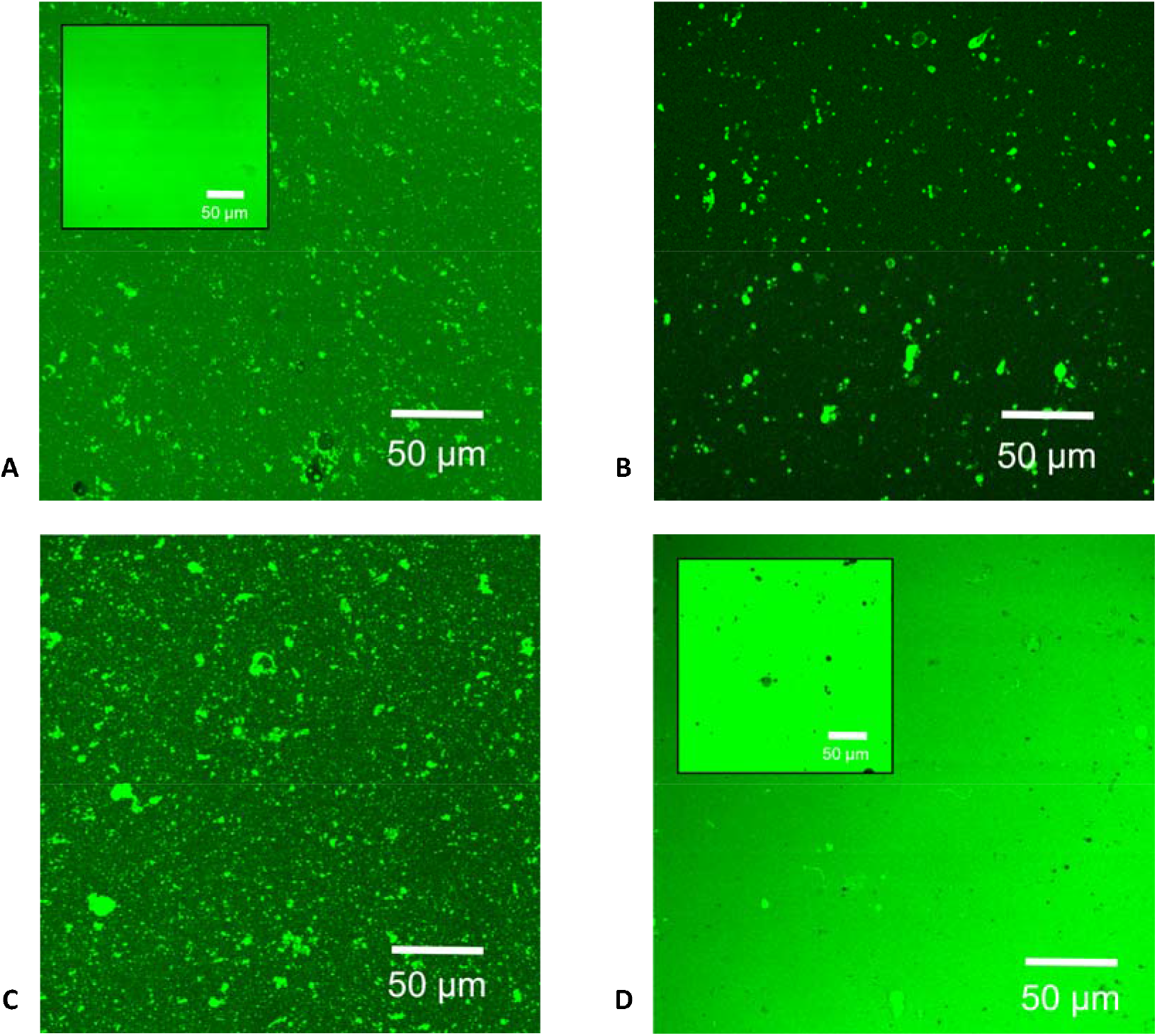
Fluorescence microscopy images of liposomes hydrated by solutions of fluorescent dyes according to Table 1: A) 5(6)-Carboxyfluorescein, B) Fluorescein, C) FITC, D) Calcein. The images show unextruded liposomes at room temperature, washed by repeated centrifugation and dilution. The insets in cases A) and D) show also “raw” liposomes before the washing steps.

On the other hand, fluorescein was visibly washed out from the liposomes during the purification procedure, which consisted of repeated centrifugation and dilution steps performed at 293 K. Ultimately, only a very small fraction of the theoretical loading capacity was retained in the liposomes (0.3 ± 0.1 μg_dye_/mg_lipid_) and there was no observable thermal release (Table 2). This is because the liposomal membrane is permeable for fluorescein even in 293 K and therefore all of F is washed out during dilution before heating. These results do not qualify fluorescein as a suitable compound for liposome formulation using the chosen lipid membrane composition due to too high permeability even at the storage temperature (logP_erm_^293 K^ = –2.6). The small amount of fluorescein that was detected after TRITON® treatment must have resided in the membrane (Fig. 4B). Such membrane partitioning is consistent with the *in silico* model, which predicted a significantly higher membrane affinity for fluorescein (logK_lip/wat_^293 K^ = 1.45) compared to 6-CF (logK_lip/wat_^293 K^ = 0.35).

The effect of high membrane partitioning coefficient was even more pronounced in the case of FITC (logK_lip/wat_^293 K^ = 2.94). Although FITC was visibly encapsulated into liposomes (Fig. 4C), no release upon heating was detected (Table 2). This was in spite of high permeability prediction for FITC, on par with fluorescein as shown in Fig. 2B. Only when a more invasive method of membrane disruption with TRITON^®^ was used, a significant release of FITC was observed (0.6 ± 0.2 μg_dye_/mg_lipid_, being 18 % of the theoretical capacity). Similarly to Fluorescein, FITC resided in the membrane rather than in the aqueous core. It can be concluded that solutes with a high partitioning coefficient toward the lipid membrane are not suitable for thermally controlled release from liposomes. This does not necessarily make lipophilic molecules unsuitable for liposomal formulations in general, but it means they are not good candidates for temperature-controlled release. Finally, let us consider Calcein. In terms of permeability predicted by the *in silio* model, Calcein was at the opposite end of the spectrum than Fluorescein or FITC, with a very low value of permeability even at elevated temperatures (logP_erm_^333K^ = –9.05) and also the lowest membrane affinity from all the investigated fluorescent dyes (logK_lip/wat_^293 K^ = −1.07). Consequently, only 0.02 % of the theoretical capacity was released after heating and this value did not substantially improve even after membrane disruption by TRITON® (Table 2). This implies that Calcein was probably not present in a majority of the liposomes already when the liposomes were formed, a hypothesis confirmed by fluorescence microscopy of “raw” liposomes observed just after the lipid hydration step (Fig. 4D, inset). The clearly visible empty liposomes surrounded by concentrated Calcein solution contrast with a similar view of 5(6)-CF liposomes (Fig. 4A, inset) where the fluorescence intensity inside and outside the raw liposomes was essentially identical.

The results of these visualisation experiments lead to a proposed mechanism of “solute rejection” during the liposome hydration step, shown schematically in Fig. 5. While well-permeating molecules such as 5(6)-CF can readily enter into the cavity of the nascent liposome, poorly permeating substances such as Calcein cannot. Therefore, the knowledge of temperature dependent membrane permeability provided by the *in silico* model is very valuable not only for predicting the liposome behaviour during storage and temperature controlled release, but also for assessing the chances that liposomes loaded by a given substance can be prepared at all.

**Figure 5.**
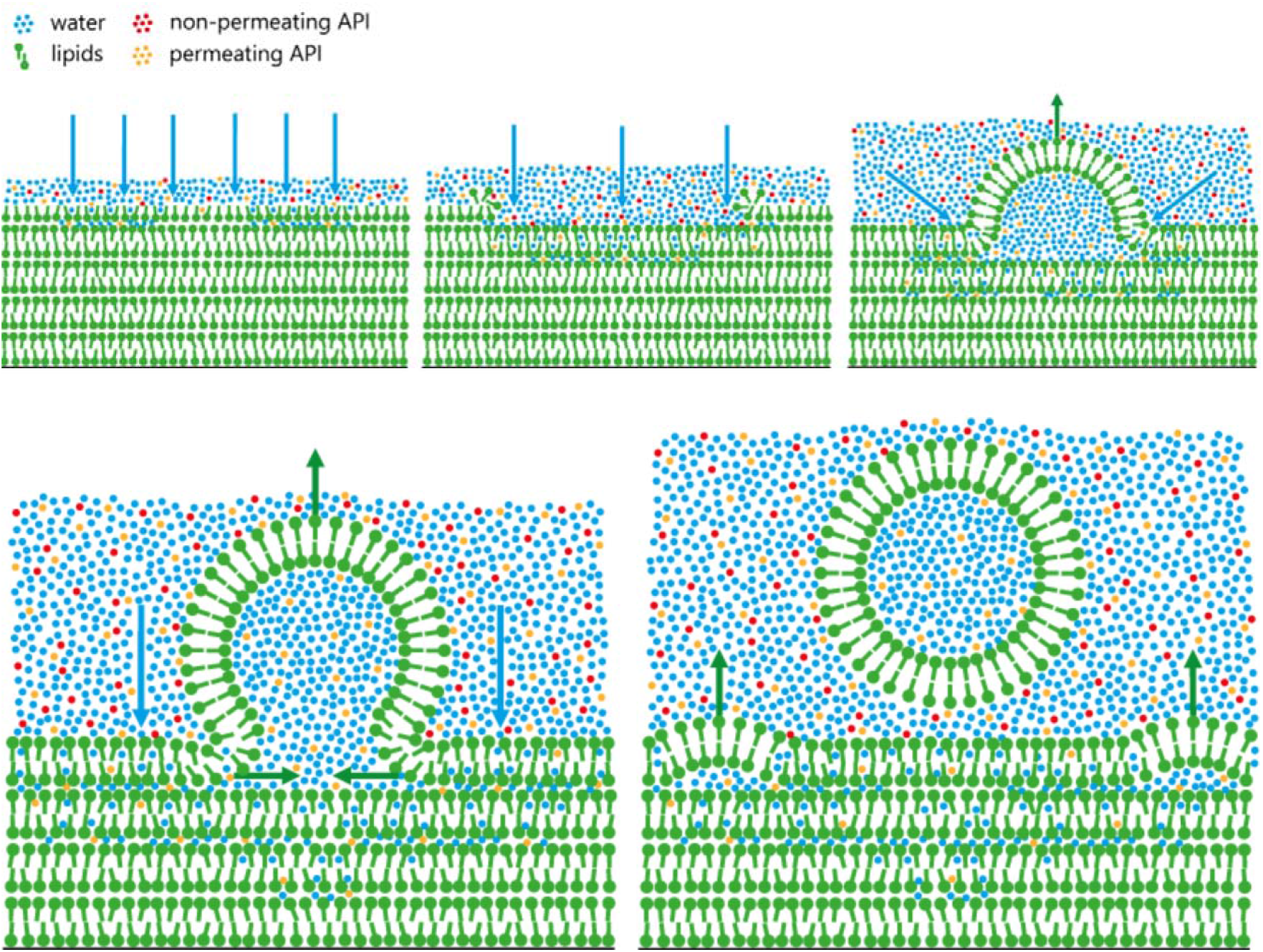
Scheme of liposome formation by the hydration of a dried lipid film. It is proposed that non-permeating solutes are excluded from the liposome cavity during this process.

### 3.3 Generalisation of rules for drug loading and thermal release from liposomes

The combination of *in silico* predictions and experimental findings discussed above, makes it possible to define a set of rules for the suitability of compounds for their incorporation into temperature controlled liposomal drug delivery systems. For each candidate molecule, the *in silico* model provides two temperature dependent coefficients: the partitioning towards the lipids (logK_lip/wat_) and the membrane permeability (logP_erm_). Given that in unilamellar liposomes, the volume of the lipidic shell is typically between 100× and 10,000× lower than the aqueous core, a logK_lip/wat_ value that ensures equal contribution of both phases to the drug loading capacity would be about 2–4 (rather than 0). This gives a limit for the encapsulation of the compound preferentially into the aqueous cavity to about logK_lip/wat_ < 2. As discussed in Section 3.2, compounds that preferentially partition into the membrane are difficult to be released thermally.

The permeability coefficient imposes two limits: *(i)* a lower limit for the storage conditions, where negligible or only very slow elution of the drug from the liposome at low temperatures is crucial to ensure stable encapsulation without premature leakage from the liposome; and *(ii)* an upper limit for the thermal release at elevated temperatures when the membrane changes to the disordered state.

To set the border values for permeation, we have to take into consideration the highest logP_erm_, where the compound did not permeate (5(6)-CF through membrane at 273 K ≈ −5.6) and the lowest logP_erm_, where the compound permeated (5(6)-CF through membrane at 333 K ≈ −3.6). We have to also taken into consideration, that only non-ionic species can permeate the membrane. Also the permeation through unstirred water should be taken into consideration as can be seen from equation 2 ^47^:

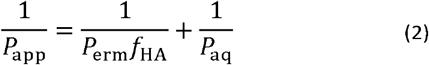

Where *P*_app_ is an apparent permeation coefficient (which is proportional to permeation time), *f*_HA_ is a fraction of the unionized species (0.0003 for 5(6)-CF at pH 7.4) and *P*_aq_ is a permeation coefficient through unstirred water layer on the membrane. The *P*_aq_ is not directly dependent on a character of permeating molecule and therefore remains constant, the apparent permeability of a compound can be estimated from calculated *P*_erm_ and *f*_HA_.

Based on the experiments and assumptions mentioned above, we propose that the ideal non-ionic compound for encapsulation and temperature-controlled release should have logP_erm_ < −9.1 at the low (ambient) temperature and logP_erm_ > −7.1 at the high (release trigger) temperature.

To set these rules more precisely, the whole procedure of liposome preparation, separation and heat release measurement was repeated with 1 mg/ml 5(6)-CF solution with PBS buffer at pH 5.4 and 6.4. The results (Table 3) shows that at lower pH, the 5(6)-CF is released at room temperature and only small amount of the dye is later released in heat induced experiment.

**Table 3.** Measured quantities of 5(6)-CF upon heat release at different pH

| 5(6)-CF<br>conc.<br>(mg/ml) | pH† | Release†† at<br>293 K | Released after<br>heating<br>($\mu\text{g}_{\text{dye}}/\text{mg}_{\text{lipid}}$) | Theoretical<br>capacity<br>($\mu\text{g}_{\text{dye}}/\text{mg}_{\text{lipid}}$) | Fraction of unionized<br>species<br>(%) | $\log P_{app}^{293K}$ |
| --- | --- | --- | --- | --- | --- | --- |
| 7.5 | 7.4 | None | $62 \pm 8$ | 246 | 0.03 % | -9.1 |
| 1 | 6.4 | Visible | $0.4 \pm 0.2$ | 32.8 | 0.26 % | -8.1 |
| 1 | 5.4 | Visible | $0.3 \pm 0.2$ | 32.8 | 2.59 % | -7.1 |

The results imply that for P_app_ = −8.1, the molecule still rapidly permeates. Since the COSMOperm method has an accuracy about 1 log value [42], more precise determination of rules for heat induced release is not necessary.

To rationalise these experiment-based rules, the calculation of time dependence of released quantity of 5(6)-CF can be made. Based on experimental approach^48^ for measurement of permeation coefficient in two compartments separated by bilayer, half of the 5(6)-CF at 293 K should be released in 2.5 hours and at 333 K or pH 5.4 at 293 K in 10 seconds (derivation in Supporting information). This is not in perfect agreement with the experiment, where not even 1 % of 5(6)-CF was released at 293 K in 25 hours [10] and therefore final rules were needed to be tuned experimentally.

According to these limits, the neutral drug molecules can be sorted into three categories:

1. Compounds not suitable for liposome encapsulation with the chosen membrane composition.
2. Compounds suitable for encapsulation but not suitable fthe thermal release mechanism.
3. Compounds suitable for both encapsulation and thermal release.

A decision tree for substance classification based on their logK__lip/wat_ and logP_erm_ values is summarised in Fig. 6A and it can be used as a basis for assessing the suitability of a given substance for incorporation into a liposomal drug delivery system.

**Figure 5.**
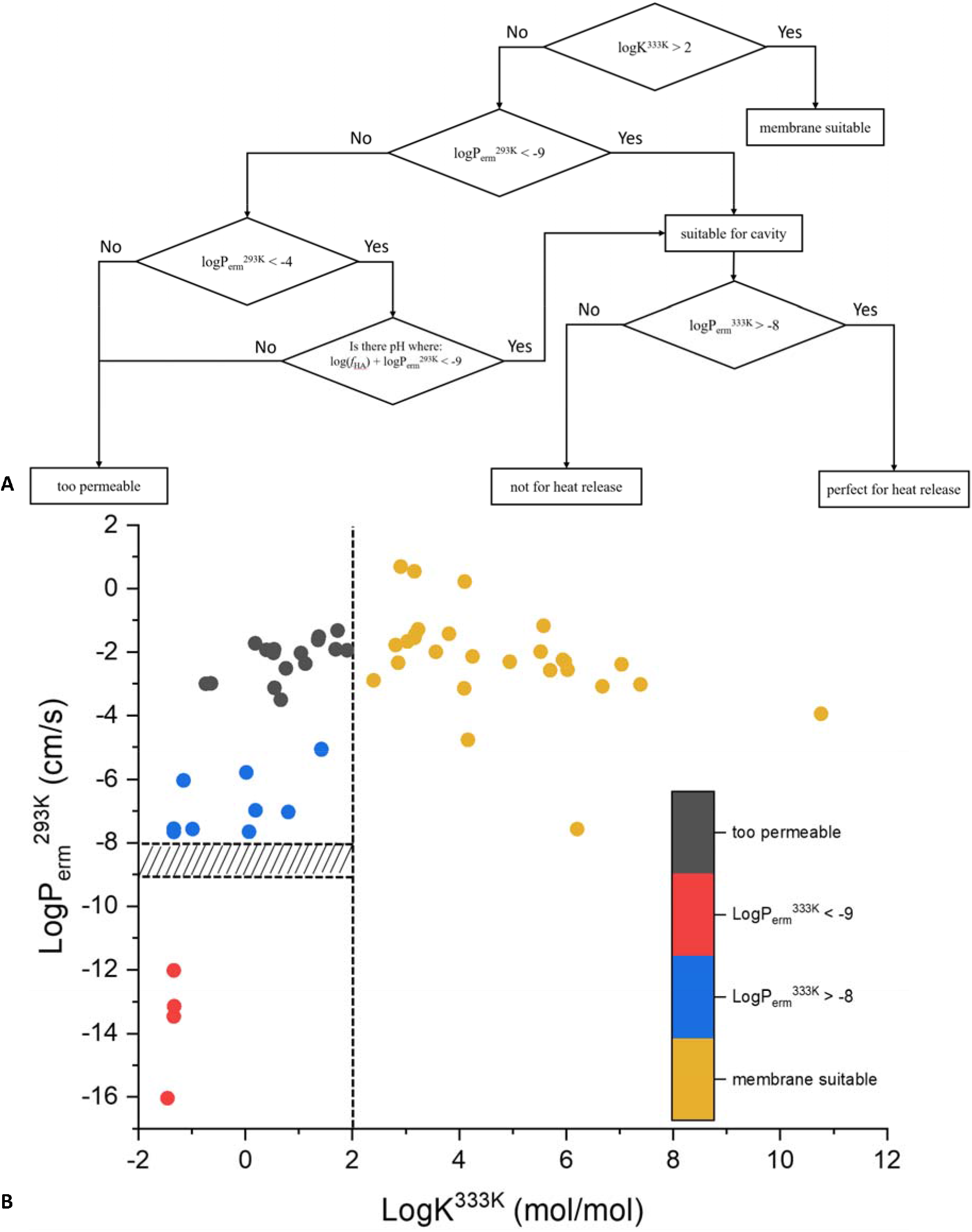
A) Decision tree dividing molecules into liposome compatibility groups according to their partition and permeation coefficients. B) Calculated partition and permeation coefficients for 56 DrugBank molecules, color-coded according to their suitability for liposome formulation (DPPC:DPPG:Chol = 75:10:15 membrane). The dotted lines denote the cut-off criteria for membrane partitioning and permeability. Molecules that are more appropriate for lipid bilayer formulation are to the right (yellow), molecules that are too permeable are in the top left corner (black), and molecules suitable for aqueous cavity formulation are further distinguished according to the likelihood of thermal release: not possible (red)and possible (blue) for which the permeability can be lowered using sufficient pH.

### 3.4 *In silico* screening of compounds from DrugBank

In order to test this set of rules, a total of 56 toxic compounds from the DrugBank database were chosen and their logK__lip/wat_ and logP_erm_ values were computed. The substances had a broad category of chemical actions including potentially harmful or deadly effect on living organisms. Since toxicity means bioactivity, such substances – even though harmful on their own – are potential candidates for applications in human or veterinary medicine, disinfection, crop protection, etc. Encapsulation into liposomes followed by controlled release could be a way how to apply such substances safely at the appropriate dose.

After applying the classification criteria (Fig. 6A) to all 56 evaluated molecules, the following groups were obtained:

- 27 substances were suitable for membrane-bound liposome encapsulation due to their high partition coefficient (yellow dots in Fig. 6B): Anacetrapib, Amitraz, Dalcetrapib, Chlorotoxin I-131, Calyculin A, Deltamethrin, Cyfluthrin, Cypermethrin, Altretamine, Bempedoicacid, Cerivastatin, Benfluorex, Coumaphos, Atorvastatin, Acifluorfen, Bisphenol A, Benzyl benzoate, Clofibrate, 1,2-dichlorobenzene, Ciprofibrate, Crotamiton, Bromoform, Bezafibrate, 9H-Carbazole, Dantron and Chlorambucil.
- 4 substances were not suitable for liposomal formulation using the Bangham method due to too low permeation even at 333 K. This could be potentially solved by using a liposome loading method that does not require membrane permeation. These compounds (red dots in Fig. 6B) were: Allosamidin, Azacitidine, Cytarabine, Decitabine,
- 16 substances were clearly not suitable for liposomal formulation due to high partitioning (black dots in Fig. 6B): Buthionine Sulfoximine, Carboquone, 2-Methoxyethanol, Acipimox, Benznidazole, Carmustine, Cerulenin, Cyclophosphamide, Carboxin, Cantharidin, Carbendazim, Cythioate, Busulfan, Allicin, Aminacrine, Chlorine
- 9 substances were identified as possible for encapsulation into the liposome aqueous cavity with a potential for thermally controlled release: Clofarabine, 8-azaguanine, Broxuridine, Cordycepin, Capecitabine, Azathioprine, Cycloserine, Dacarbazine, PhIP.
- The details of 9 compounds identified as potentially suitable for liposome encapsulation and thermal release are summarised in Table 4. The pH range for fHA needed was calculated using Protonation Plugin Group in Marvin Sketch 20.16 [44]. All of the compound has a theoretical window of pH opportunity to be encapsulated into liposomes and later thermally released. On the other hand, some of the theoretically viable pH ranges are strongly acidic or basic and therefore practical encapsulation would be very problematic.

**Table 3.** Calculated logP_erm_ and logK values for 9 toxic compounds in neutral form that were identified as possible candidates for encapsulation and thermal release from DPPC:DPPG:Chol (75:10:15) liposomes.

| compound | $\log K^{293K}$ | $\log P_{\text{erm}}^{293K}$<br>(cm/s) | $\log K^{333K}$ | $\log P_{\text{erm}}^{333K}$<br>(cm/s) | $f_{\text{HA}}$ needed<br>(%) | pH range |
| --- | --- | --- | --- | --- | --- | --- |
| Clofarabine | -1.3 | -7.66 | -1.34 | -5.44 | 4.6 | 0 – 0.8 |
| 8-azaguanine | 0.71 | -7.65 | 0.07 | -5.85 | 4.5 | 7.2 – 14 |
| Broxuridine | -1.3 | -7.57 | -0.99 | -5.54 | 3.7 | 9.8 – 14 |
| Cordycepin | -1.31 | -7.55 | -1.34 | -5.44 | 3.5 | 0 – 2.6 |
| Capecitabine | 0.02 | -7.03 | 0.8 | -4.71 | 1.1 | 10.8 – 14 |
| Azathioprine | 0.54 | -6.98 | 0.19 | -5.18 | 0.095 | 0 – 0.6,<br>10.8 – 14 |
| Cycloserine | -0.83 | -6.04 | -1.15 | -4.6 | 0.11 | 0 – 14* |
| Dacarbazine | 0.4 | -5.78 | 0.02 | -4.15 | 0.058 | 12.2 – 14 |
| PhIP | 2.56 | -5.06 | 1.42 | -3.88 | 0.011 | 0 – 0.8 |

\*Cycloserine is present mainly in zwitterionic and not neutral form.

### 3.5 Liposomal formulation of cycloserine delivery

As a final experimental validation of the *in silico* screening methodology, one of the most promising candidates was selected and encapsulated into liposomes. Since D-cycloserine is reasonably safe to work with and has well-documented antibiotic properties, this compound was chosen and tested in experiments with bacteria (*E. coli*) as described in Section 2.3. The roughly estimated *P*_app_ (Supporting information S5) at pH 7.4 is −10.1 at 293 K and −8.66 at 333 K. This means that cycloserine is clearly not permeating at low temperature but can be permeating at higher temperature because the permeation coefficient is at uncertain region between −9 and −8. The results of the resazurin assay are summarised in Figure 7. The ability of the liposomes to retain the encapsulated bioactive compound cycloserine was evidenced by a control group exposed to the supernatant from intact liposomes, whose resazurin metabolic activity was almost the same as that of the negative control (pure bacteria). On the other hand, bacteria groups exposed to cycloserine that was released from liposomes after heat or TRITON® treatment exhibited a substantially reduced metabolic activity. The antibacterial effect was not as strong as that of the control group exposed to 50 μg/mL kanamycin, but that is only a matter of the applied dose. Crucially, there was no statistical difference between the antibacterial effect of TRITON^®^ release and heat release groups, which means that cycloserine is indeed a suitable compound for liposomal formulation and release by the heating mechanism.

**Figure 7.**
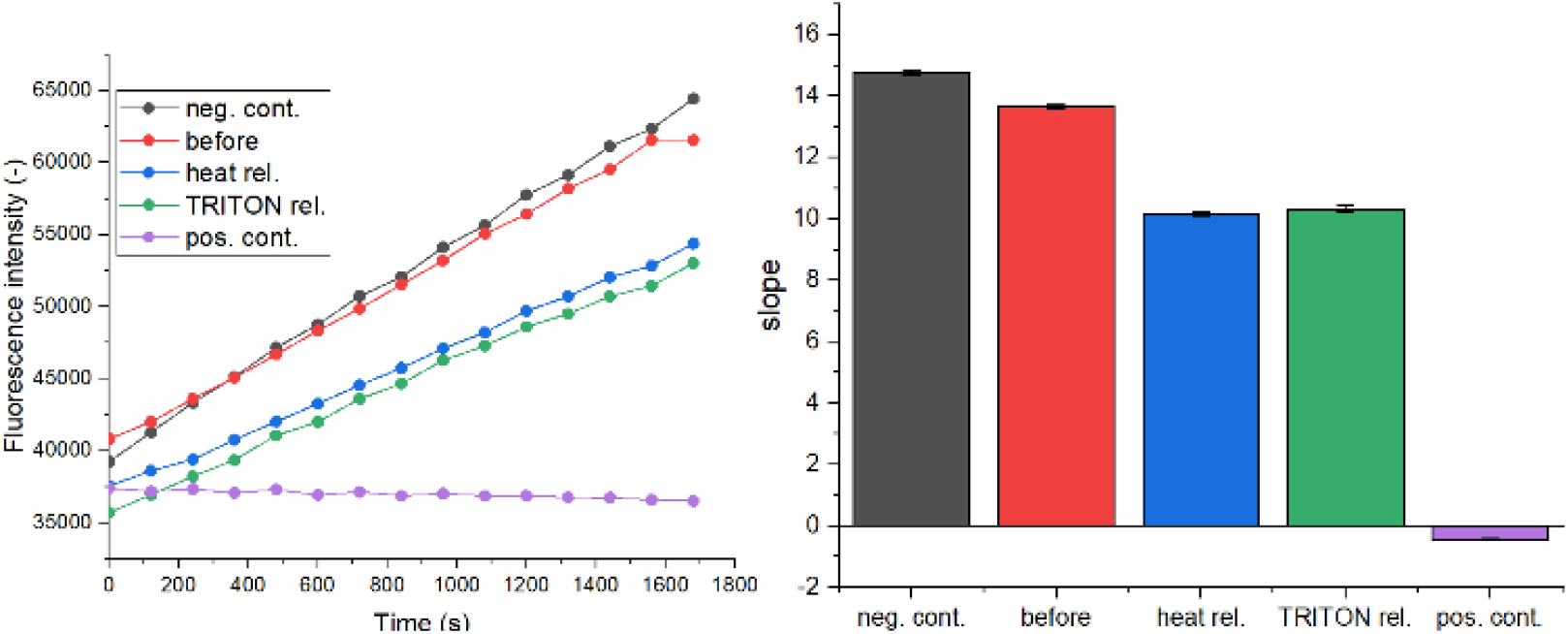
Effect of cycloserine on E. coli tested by the Resazurin assay. A) Dependence of measured fluorescence intensity on time (each line was averaged from 3–6 wells). B) Calculated slopes from the fluorescence measurement for individual control groups. Neg. cont. – bacteria only; before – bacteria exposed to supernatant from cycloserine loaded liposomes kept at ambient temperature; heat. rel. – bacteria exposed to supernatant from cycloserine loaded liposomes after heat release; TRITON rel. – bacteria exposed to supernatant from cycloserine loaded liposomes after membrane disruption by TRITON®; pos. cont. – bacteria exposed to 50 μg/mL kanamycin.

In result, our *in silico* methodology benchmarked on fluorescein derivatives was successful in the selection of cycloserine as a valid candidate for liposomal formulation with heat release and selected lipid composition.

## 4 Conclusions

A new *in silico* methodology for the selection of potent candidate substances suitable for the encapsulation and thermally controlled release from liposomes has been proposed and verified experimentally. The whole procedure comprises molecular dynamic simulation of small patch of a bilayer with defined composition used for liposome formulation, COSMOperm/COSMOmic calculations for candidate molecules in neutral form and their selection using a set of experimentally validated rules. In this work, we used bilayer composition DPPC:DPPG:Chol (75:10:15 molar ratio) and 56 drug candidates. Comparison between experimentally measured thermal release and COSMOperm/COSMOmic calculations of a set of fluorescent dyes enabled us to derive rules for the selection of molecules that can be encapsulated in liposomal cavities and thermally released. These rules were then applied for the prediction of liposomal compatibility of 56 toxic compounds from the DrugBank database. From this set of compounds, 16 showed no suitability for liposome encapsulation, 27 showed suitability for encapsulation into the liposome membrane, 4 showed suitability for cavity encapsulation using methods that do not involve membrane permeation during liposomes loading, and finally 9 compounds were deemed suitable for encapsulation into the liposome cavity. One of these compounds – cycloserine – was then selected for a successful experimental demonstration of liposome encapsulation and thermal release.

In the present work, the composition of the lipid bilayer and the structure of the tested molecules were fixed. However, both represent additional potential degrees of freedom. For example, the *in silico* methodology developed in this work can be extended for finding such combination of lipids in the liposome membrane that will meet the logK and logP_erm_ criteria for a specific molecule whose liposome encapsulation is highly desirable. Such computational formulation optimisation can save valuable experimental resources and lead to novel compositions that might not be found experimentally. Similarly, during the development of new pharmaceutically active compounds, the lead structure optimisation can be partially driven by criteria for logK and logP_erm_ that were proposed and validated in the present work. In this way, new chemical entities can be optimised not only for their biological efficacy, but also for formulation manufacturability.

## Author contribution

M.B. – liposome loading/release experiments, TEM measurements, data analysis, manuscript writing

M.Šr. – computer simulations, data analysis

M.Šo. – liposome loading/release experiments, data analysis, confocal analysis

P. J. – bacterial assay experiments, data analysis

F.Š. – conceptualisation, data interpretation, manuscript writing, funding acquisition

K.B. – conceptualisation, computer simulations, data analysis, manuscript writing, funding acquisition

## Acknowledgements

M.B., M.Šo. and P.J. acknowledge support by Specific University Research (project MŠMT no. 21-SVV/2020). M.Šr. acknowledges supported by Palacký University Olomouc (project IGA_PrF_2020_022). F.Š. would like to thank the Czech Science Foundation (project no. 19-09600S) for financial support. K.B. and M.Šr. would like to thank the Czech Science Foundation (project no. 17-21122S).

