## Supplemental Information for "In-silico screening of drug candidates for thermoresponsive liposome formulations"

S1: Analysis of molecular dynamic simulations

To evaluate if the membrane in Molecular dynamic (MD) simulation is already equilibrated, analysis of area corresponding to one lipid was performed (Fig. S1.1). It was observed that areas per lipid in a case of MD simulation at 293 K, 313 K and 323 K did not vary a lot although it can be seen slight increase with the temperature. On the other hand, MD simulation at 333 K showed rapid increase in area per lipid. In all cases, there is no systematic change in area per lipid after 100 ns and therefore time of equilibration around 200 ns is definitely sufficient for our simulation.

 
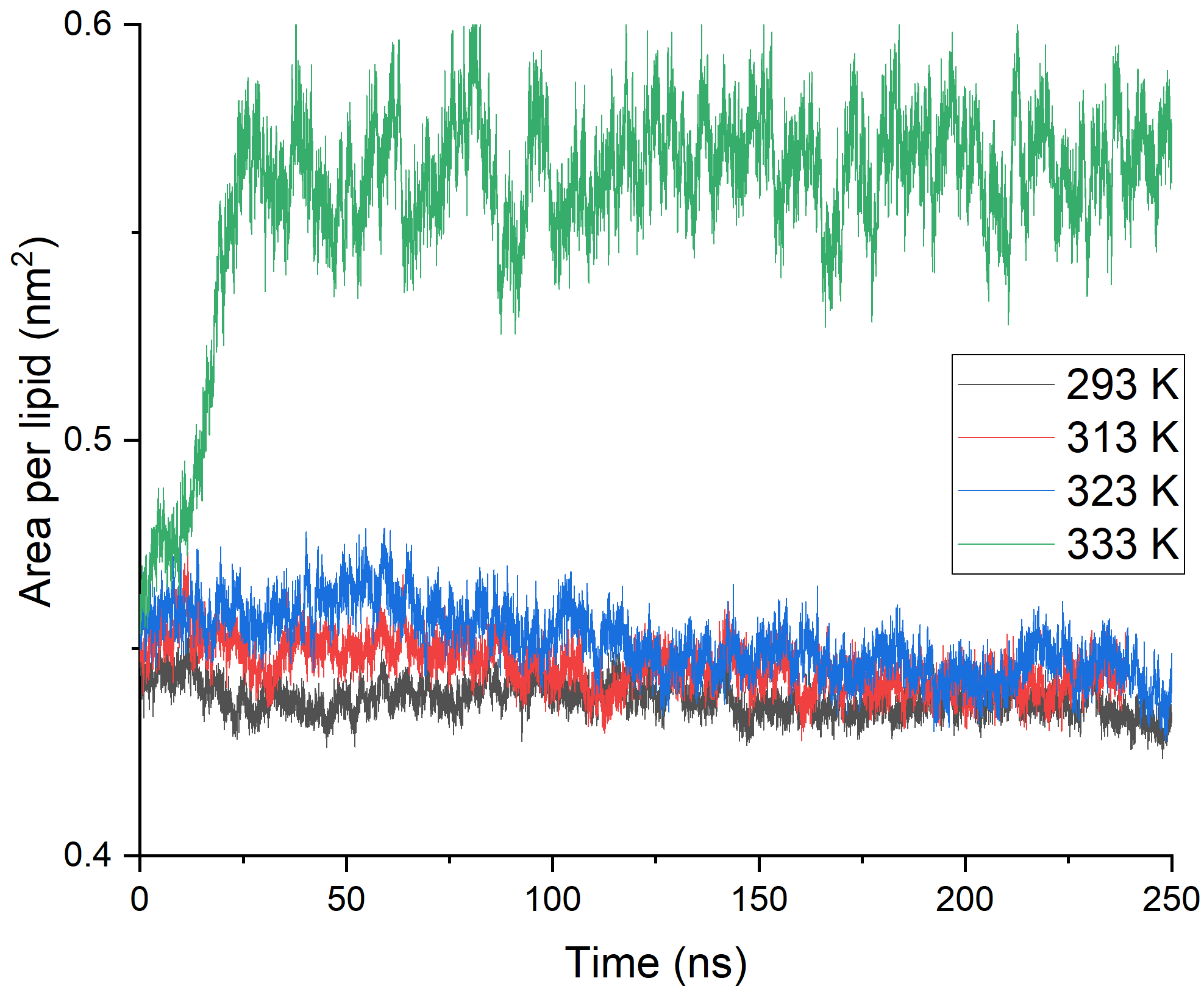

**Figure S1.1:** Dependency of area per lipid on a simulation time for 4 different temperatures of simulated membrane DPPC:DPPG:Chol (75:10:15)

Although the different behavior of membrane at 333 K can be clearly seen from above, order parameters [1] for both (sn1 and sn2) palmitates in DPPC were calculated using *gmx order* [2] tool (Fig. S1.2). The disorder of a membrane at 333 K is then clearly seen from the comparison with MD simulations at lower temperatures.

 
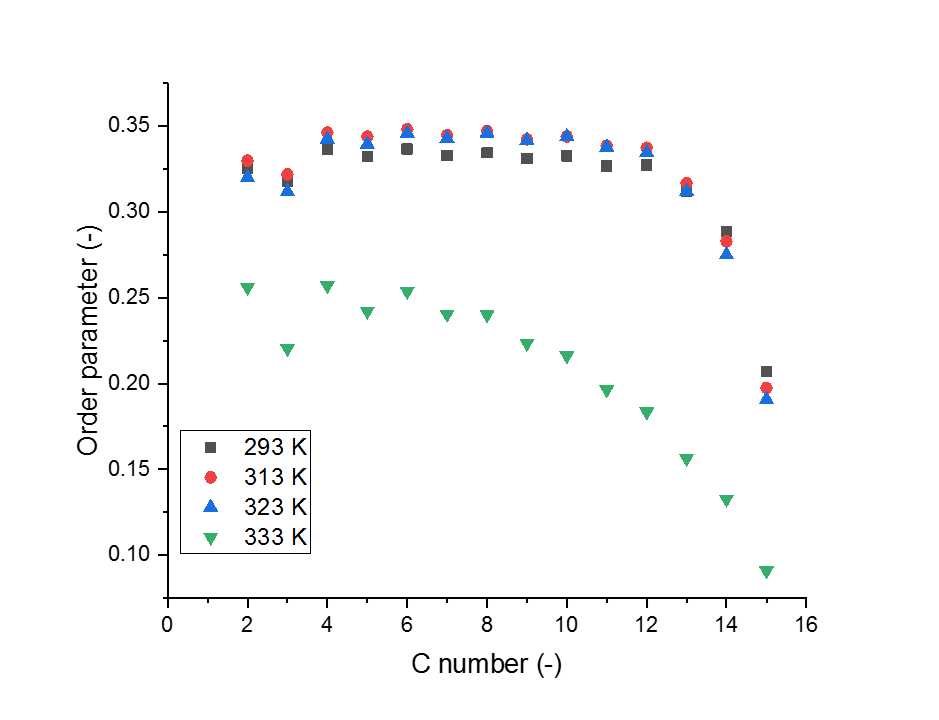
 
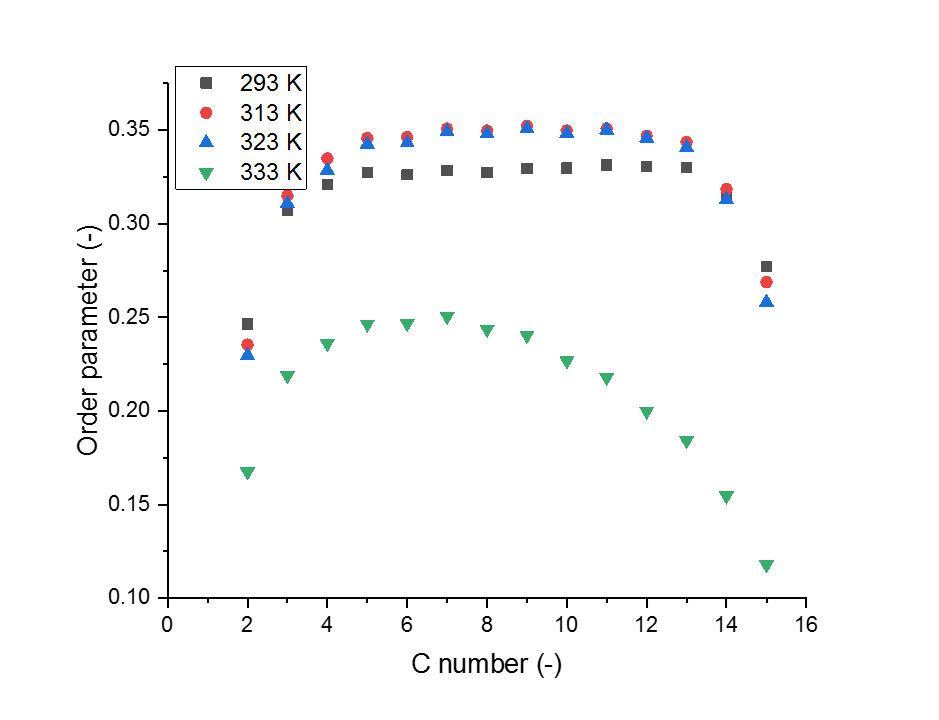

**Figure S1.2:** Order parameters for sn1 (left) and sn2 (right) palmitates in DPPC

S2: Calculated energy profiles through membrane

The calculation of partition and permeation coefficient is based on COSMOmic approach which is described in detail in COSMOmic publication [3]. Here briefly:

From the equilibrated MD simulation of lipid bilayer, a screenshot is taken and a bilayer was cut to 50 slices along the *r* axis that goes through membrane (Fig 2.1).

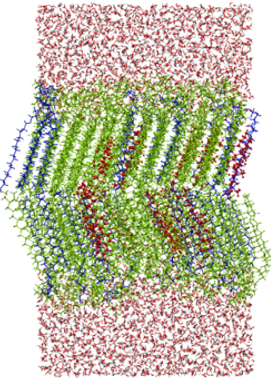

*r*

**Figure S2.1:** Screenshot from MD simulation used for partitioning and permeation calculation with the indication of slices which are used for energy profile calculation.

For each slice, σ-profile is calculated and using the COSMO-RS approach and σ-profile of a permeating molecule, chemical potential of a molecule (i) in each layer is calculated as a function of orientation in space of the molecule ($\mu_{i}(r,d))$.

From the chemical potential, partition function in each slice can be calculated from chemical potentials of all orientations taken into consideration:

$$Z_{i}\left( r \right)=\sum_{d} e^{\frac{-\mu_{i}(r,d)}{kT}}$$

(S1)

From the known partition function of molecule in last layer where only water is present ($Z_{i}\left( n \right)$), the relative partitioning and therefore also free energy difference to water environment can be calculated:

$${\Delta G}_{i}\left( r \right)=-RT\ln\frac{Z_{i}\left( r \right)}{Z_{i}\left( n \right)}$$

(S2)

The difference in free energy can be taken into graph and evaluated as a function of the distance from membrane center (Fig. S2.2). Moreover, from the free energy difference in a local minimum along r axis, the partition coeficient logK_lip/wat_ is calculated:

$$\log K_{lip/wat}=e^{\frac{-{\Delta G}_{i}\left( \min\right)}{RT}}$$

(S3)

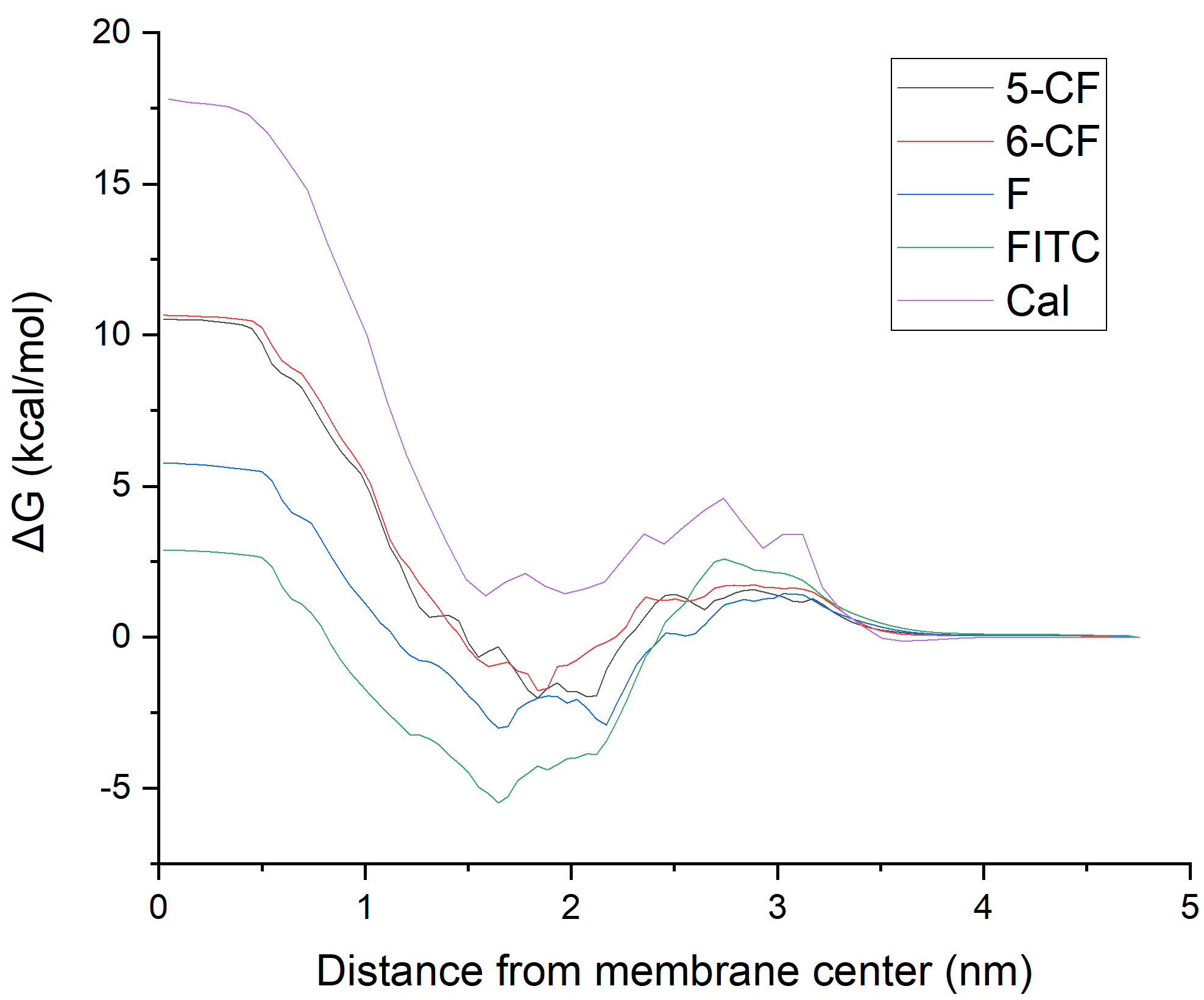

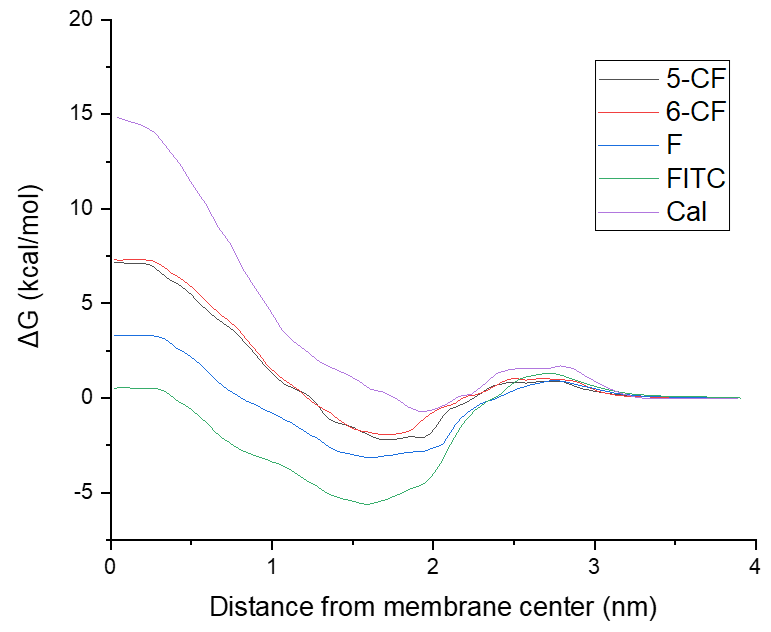

**Figure S2.2:** Calculated energy profiles through DPPC:DPPG:Chol (75:10:15) membrane for 4 fluorescein derivatives (5-CF and 6-CF as a mixture in experiments) at 293 K (left) and 333 K (right) used as a calibration of whole procedure.

S3: Calculation of logP_erm_

Calculation of permeation coefficient is explained in detail in COSMOperm publication [4] and is based on a equation firstly proposed by Diamond and Katz [5]:

$$\frac{1}{P_{\mathrm{erm}}}=\int_{-L}^{L} \frac{1}{K\left( z \right)D(z)}dz$$

(S4)

where $L$ is thickness of one lipid layer, $K(z)$ is partition coefficient in each slice and $D(z)$ is diffusion coefficient in each slice. Partition coefficient is calculated from the change of free energy using the procedure described in section S2.

Diffusion coefficient is calculated using COSMO-RS approach using experimentally fitted parameters and COSMO profile of a permeating molecule.

S4: Racionalisation of time of permeability calculation

From the experimentally used procedure for calculation of apparent permeation coefficient (*P*_app_) [6]:

$$P_{\mathrm{app}}=slope\cdot\frac{V_{\mathrm{cell}}}{S_{m}\cdot c_{\mathrm{donor}}}$$

(S5)

where $slope$ is the slope in dependency of acceptor concentration on time, $V_{\mathrm{cell}}$ is the volume of cell, $S_{m}$ is the area of permeating membrane and $c_{\mathrm{donor}}$ is concentration in donor phase.

If we take into consideration that we do not have 2 equally big compartments but many liposomes with dye and one “big” acceptor compartment, the equation can be rewritten into new form:

$$P_{\mathrm{app}}=\frac{{dc}_{\mathrm{acc}}}{dt}\cdot\frac{V_{\mathrm{acc}}}{S_{\mathrm{lip}}\cdot N_{\mathrm{lip}}\cdot c_{\mathrm{lip}}(t)}$$

(S6)

where $\frac{{dc}_{\mathrm{acc}}}{dt}$ is the time derivation of acceptor concentration, $V_{\mathrm{acc}}$ is the volume of acceptor compartment (1.6 ml in our experiments), $S_{\mathrm{lip}}$ is the area of one lipid, $N_{\mathrm{lip}}$ is the number of lipids in the sample and $c_{\mathrm{lip}}(t)$ is the concentration of the dye in liposomes but i tis time dependent as the concentration of dye in liposome during permeation is changing rapidly.

The whole amount of dye which comes to the permeation experiment is on the beginning of the experiment in the liposomes with starting concentration $c_{0}$ and in each time, the dye is either in liposomes or in akceptor compartment.

$$c_{0}V_{\mathrm{lip}}N_{\mathrm{lip}}=c_{\mathrm{lip}}V_{\mathrm{lip}}N_{\mathrm{lip}}+c_{\mathrm{acc}}V_{\mathrm{acc}}$$

(S7)

The dependency of concentration in acceptor phase on the concentration in lipids:

$$c_{\mathrm{acc}}=\frac{V_{\mathrm{lip}}N_{\mathrm{lip}}}{V_{\mathrm{acc}}}\left( c_{0}-c_{\mathrm{lip}} \right)$$

(S8)

And the derivation:

$$\frac{{dc}_{\mathrm{acc}}}{{dc}_{\mathrm{lip}}}=-\frac{V_{\mathrm{lip}}N_{\mathrm{lip}}}{V_{\mathrm{acc}}}$$

(S9)

The derivation in (S6) can be rewritten:

$$\frac{{dc}_{\mathrm{acc}}}{dt}=\frac{{dc}_{\mathrm{acc}}}{{dc}_{\mathrm{lip}}}\frac{{dc}_{\mathrm{lip}}}{dt}$$

(S10)

Using (S10) and (S9), the (S6) can be written in a form, where only time-dependent variable is concentration in liposomes.

$$P_{\mathrm{app}}=-\frac{V_{\mathrm{lip}}N_{\mathrm{lip}}}{V_{\mathrm{acc}}}\frac{{dc}_{\mathrm{lip}}}{dt}\cdot\frac{V_{\mathrm{acc}}}{S_{\mathrm{lip}}\cdot N_{\mathrm{lip}}\cdot c_{\mathrm{lip}}}$$

(S11)

$$P_{\mathrm{app}}=-\frac{{dc}_{\mathrm{lip}}}{dt}\cdot\frac{V_{\mathrm{lip}}}{S_{\mathrm{lip}}\cdot c_{\mathrm{lip}}}$$

(S12)

With an assumption that liposomes are spherical with diameter (*d*) equal to 600 nm:

$$P_{\mathrm{app}}=-\frac{{dc}_{\mathrm{lip}}}{dt}\cdot\frac{d}{6\cdot c_{\mathrm{lip}}}$$

(S13)

The differential equation has with initial condition ($c_{\mathrm{lip}}\left( 0 \right)=c_{0})$solution:

$$\ln c_{\mathrm{lip}}=\ln c_{0}-\frac{6\cdot P_{\mathrm{app}}}{d}t$$

(S14)

If we choose some characteristic release (half time of release), we get equation for half-time:

$$t_{1/2}=\frac{d\cdot\ln2}{6\cdot P_{\mathrm{app}}}$$

(S15)

For the $P_{\mathrm{app}}$ of 5(6)-CF at 293 K (${10}^{-7.1}$) we get half-time 87 seconds and for 5(6)-CF at 333 K (${10}^{-9.1}$) we get 2.4 hours.

S5: Calculated values for drug candidates

For a total number of 57 compound, partition and permeation coefficients at 293 K and 333 K were calculated. All calculated values are listed in Table S4.1.

**Table S4.1:** COSMOmic calculated partition coefficients and COSMOperm caclculated permeation

| compound | logK^293K^ | logP_erm_^293K^ (cm/s) | logK^333K^ | logP_erm_^333K^ (cm/s) |
| --- | --- | --- | --- | --- |
| 1_2-dichlorobenzene | 3.28 | -1.29 | 3.23 | 0.06 |
| PhiP | 2.56 | -5.06 | 1.42 | -3.88 |
| 2-Methoxyethanol | -1.21 | -3.00 | -0.74 | -1.58 |
| 8-azaguanine | 0.71 | -7.65 | 0.07 | -5.85 |
| 9H-CARBAZOLE | 3.23 | 0.69 | 2.91 | 0.32 |
| Acifluorfen | 4.59 | 0.22 | 4.10 | 1.09 |
| Acipimox | -0.86 | -2.99 | -0.64 | -1.87 |
| Allicin | 1.28 | -1.62 | 1.36 | -0.54 |
| Allosamidin | -1.47 | -16.04 | -1.45 | -12.09 |
| Altretamine | 5.41 | -2.58 | 5.70 | -0.06 |
| Aminacrine | 1.29 | -1.52 | 1.38 | -0.47 |
| Amitraz | 7.07 | -3.03 | 7.39 | -0.12 |
| Anacetrapib | 10.40 | -3.95 | 10.77 | -0.36 |
| Atorvastatin | 4.02 | -4.77 | 4.16 | -1.41 |
| Azacitidine | -1.23 | -13.46 | -1.34 | -10.56 |
| Azathioprine | 0.54 | -6.98 | 0.19 | -5.18 |
| Bempedoic acid | 4.81 | -1.17 | 5.58 | 0.69 |
| Bendamustine | 2.87 | -2.89 | 2.40 | -0.92 |
| Benfluorex | 7.97 | -2.31 | 4.94 | -0.12 |
| Benznidazole | 1.07 | -2.52 | 0.76 | -1.44 |
| Benzylbenzoate | 3.89 | -1.43 | 3.81 | 0.05 |
| Bezafibrate | 3.11 | -1.67 | 3.03 | 0.00 |
| Bisphenol A | 4.16 | -3.15 | 4.09 | -0.82 |
| Bromoform | 2.94 | 0.54 | 3.16 | 0.74 |
| Broxuridine | -1.30 | -7.57 | -0.99 | -5.54 |
| Busulfan | 0.48 | -1.73 | 0.18 | -1.11 |
| Buthionine Sulfoximine | 0.83 | -3.51 | 0.66 | -2.30 |
| Calyculin A | 4.95 | -7.57 | 6.20 | -2.49 |
| Cantharidin | 0.22 | -1.93 | 0.40 | -0.82 |
| Capecitabine | 0.02 | -7.03 | 0.80 | -4.71 |
| Carbendazim | 0.63 | -1.92 | 0.54 | -0.97 |
| Carboquone | 0.46 | -3.13 | 0.55 | -1.41 |
| Carboxin | 1.93 | -1.94 | 1.91 | -0.68 |
| Carmustine | 1.09 | -2.37 | 1.12 | -0.92 |
| Cerivastatin | 5.13 | -1.99 | 5.52 | 0.28 |
| Cerulenin | 0.51 | -2.04 | 1.04 | -1.10 |
| Chlorambucil | 2.99 | -1.78 | 2.81 | -0.17 |
| Chlorine | 1.58 | -1.33 | 1.72 | -0.17 |
| Chlorotoxin I-131 | 7.07 | -3.08 | 6.68 | -0.60 |
| Ciprofibrate | 3.34 | -1.44 | 3.18 | 0.19 |
| Clofarabine | -1.30 | -7.66 | -1.34 | -5.44 |
| Clofibrate | 3.46 | -2.00 | 3.56 | -0.36 |
| Cordycepin | -1.31 | -7.55 | -1.34 | -5.44 |
| Coumaphos | 4.20 | -2.14 | 4.25 | -0.12 |
| Crotamiton | 2.89 | -1.55 | 3.16 | 0.10 |
| Cyclophosphamide | 0.28 | -2.03 | 0.53 | -0.96 |
| Cycloserine | -0.83 | -6.04 | -1.15 | -4.60 |
| Cyfluthrin | 6.07 | -2.30 | 5.97 | -0.16 |
| Cypermethrin | 6.03 | -2.25 | 5.94 | -0.13 |
| Cytarabine | -0.82 | -13.14 | -1.33 | -10.30 |
| Cythioate | 2.47 | -1.92 | 1.69 | -0.93 |
| Dacarbazine | 0.40 | -5.78 | 0.02 | -4.15 |
| Dalcetrapib | 6.65 | -2.39 | 7.03 | -0.01 |
| Dantron | 2.94 | -2.34 | 2.86 | -0.62 |
| Decitabine | -1.26 | -12.03 | -1.34 | -9.34 |
| Deltamethrin | 6.23 | -2.56 | 6.03 | -0.33 |

S6: Calculation of *f*_HA_ for cycloserine

Using the Protonation Plugin Group in MarvinSketch 20.16 [7], and the pK_A_ for the deprotonation of secondary amine is 4.21 (pK_A1_) and the pK_A_ for protonation of primary amine is 8.34 (pK_A2_). Then, 4 forms of cycloserine can be present in solution (Fig. S6.1).

From the equilibria, the equation for ratio between neutral and zwitterionic form can be written.

$$\frac{c_{N}}{c_{\mathrm{Zw}}}=\frac{K_{A2}}{K_{A1}}={10}^{{pK}_{A1}-{pK}_{A2}}=7.4\cdot{10}^{-5}$$

(S16)

This ratio is based on rough theoretical prediction of acidity constants and therefore can vary from the reality.

The zwitterionic form is present at pH 7.4 in 90 % and therefore $f_{HA}$ for neutral cycloserine at this pH is $6.67\cdot{10}^{-5}$. Finally, the apparent permeation coefficient is -10.2 and -8.8 at 293 K and 333 K, respectively.

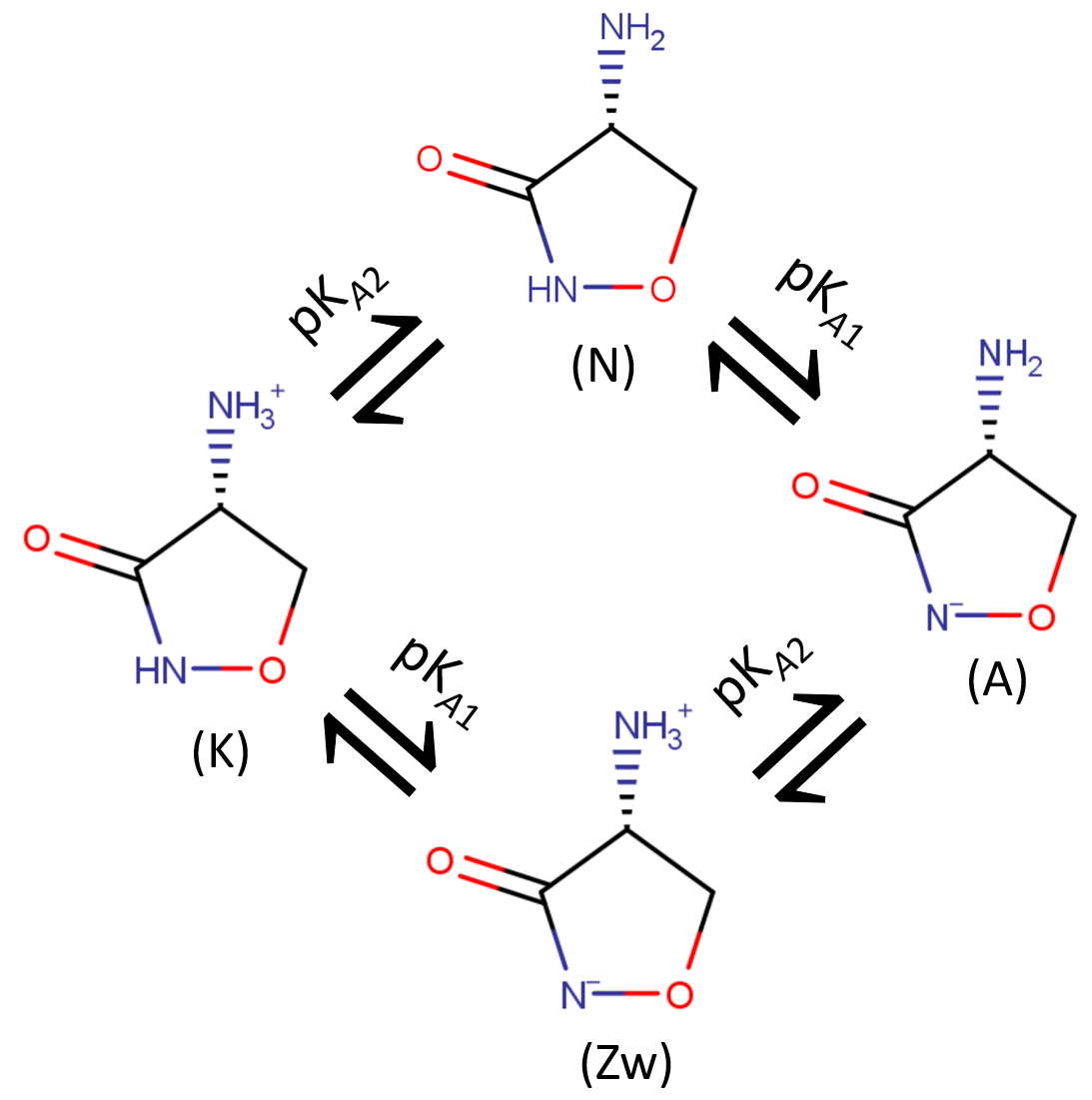

**Figure S6.1:** Forms of cycloserine and their equilibria in water solution: kationic (K), neutral (N), anionic (A) and zwitterionic (Zw)

7.	*MarvinSketch 20.16*. 2020, ChemAxon Ltd.
